# Large-scale quantum computing framework enhances drug discovery in multiple stages

**DOI:** 10.64898/2026.02.09.704961

**Authors:** Kai Wen, Jinyin Zha, Shaobo Chen, Jie Zhong, Lixin Yuan, Yunxia Cui, Xinchao Shi, Wenming Qin, Xiaobing Lan, Yonghui Liu, Xiuyan Yang, Hui Qin, Mingyu Li, Putuo Guo, Qingjie Xiao, Tingting Wu, Yuhui Zhou, Chongyu Cao, Shaobo Ning, Chengwei Wu, Qi Gao, Hongxi He, Yao Ma, Zhenyu An, Xinyi Liu, Yingyi Chen, Zhen Zheng, Hai Wei, Yin Ma, Jian Zhang

## Abstract

Coherent Ising machines (CIMs) excel at solving large-scale combinational optimization problems (COPs), but their insufficient long-term stability has hindered their applications in compute-intensive tasks like computer-aided drug discovery (CADD). By improving fiber vibration isolation and temperature control system, we have implemented a 2000-node CIM named *QBoson-CPQC-3Gen* achieving stable solutions over one hour on large-scale COPs. Graph-based encoding schemes were further introduced to realize a CIM-based CADD workflow including allosteric site detection, protein-peptide docking and intermolecular similarity calculation. CIM-based methods demonstrated superior speed and accuracy than heuristic algorithms. Especially, *QBoson-CPQC-3Gen* identified 2 novel druggable sites and bioactive compounds for 6 targets, which were further validated *in vitro*, in-cell and by crystal structures. Our contributions established a quantum-computing framework for multi-stage drug discovery, representing a significant advancement in both quantum computing applications and pharmaceutical research.

## Introduction

Drug discovery^1–3^ typically involves significant investments of time and resources. Computer-aided drug discovery (CADD)^4–6^ is widely believed to enhance efficiency and reduce associated costs in this field. However, the frequent emergence of large-scale combinatorial optimization problems^7^ (COP) within CADD poses considerable challenges to both computational speed and accuracy. COPs are optimization problems defined over discrete solution spaces and are classified as NP-hard; as a result, they are primarily addressed using heuristic algorithms on digital computers^8–10^. These algorithms often face a trade-off between solution quality^11^ and time efficiency^12^. To overcome these limitations, various quantum computing platforms like quantum annealers^13^, gaussian boson samplers^14^ and coherent Ising machines^15–17^ (CIMs) have been developed, which tackle COPs by leveraging the evolution of qubit systems. Among these, CIMs are notably capable of accommodating larger numbers of fully connected spins^18–20^, making them particularly suitable for large-scale COPs in CADD.

Nevertheless, the current generation of CIMs faces challenges in advancing CADD applications. On one hand, many CADD necessitate repeated calculations of large-scale COPs^6^, demanding both high qubit counts and solution stability. Although dense-qubit CIMs have been reported^18^, these systems often exhibit a rapid decline in solution quality over time^20^. On the other hand, encoding diverse COPs into CIM-readable matrices remains a methodological prerequisite, and the exploration of encoding strategies specific to CADD-related COPs is still limited^21–24^. Especially, there is currently no end-to-end, CIM-based framework capable of initiating from a disease protein and proceeding through a full drug-design pipeline. Moreover, benchmarking in existing studies has largely been confined to individual cases^24, 25^.

Here, we report a stable coherent Ising machine named QBoson-CPQC-3Gen that can solve COPs up to 2000 binary variables. Through improved fiber thermostat control and enhanced vibrational isolation of the optical path, this system achieves continuous and stable operation on large-scale COPs for over one hour at room temperature. To leverage this robust platform, we developed graph-based encoding schemes that integrate *allosteric site prediction*, *protein-peptide docking,* and *intermolecular similarity calculation* into a unified CIM framework, which collectively enable prediction of druggable regions disease proteins, identification of active compounds binding to those regions, and the expansion of active compound libraries, forming an end-to-end, CIM-powered pipeline for drug discovery. We extensively benchmarked the performance of these algorithms across multiple datasets. Importantly, through combined CIM computations and wet-lab validations (Table 1), we successfully identified two previously unreported allosteric sites, two novel peptide modulators, and several small-molecule modulators. These results underscore the pioneering potential of CIMs in real-world drug discovery applications.

**Table 1.**
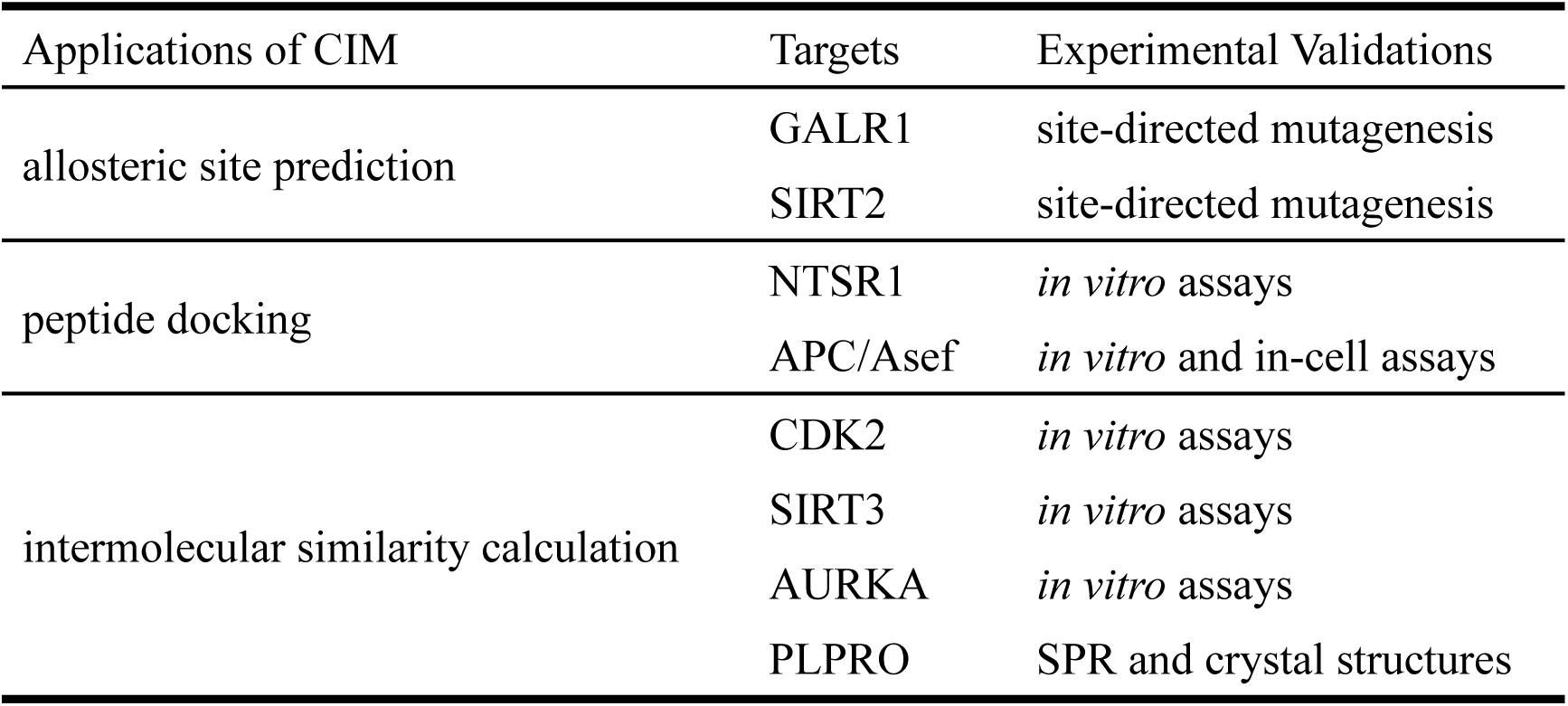
List of experimental validation targets.

## Results

### QBoson-CPQC-3Gen is a stable CIM for large-scale COPs

In a coherent Ising machine, degenerate optical parametric oscillators (DOPOs), generated within a nonlinear optical crystal, function as artificial spins. A DOPO assumes a phase of either 0 or π relative to the pump phase when operating above threshold. Maintaining stable DOPO phases is crucial for achieving high solution quality with CIM. However, earlier CIM implementations have struggled to sustain DOPO stability over extended durations, particularly when solving large-scale problems^20^. even though recent advances have enabled CIMs to support over 100,000 spins^18^, their use in practical settings that require high-frequency computation remains limited.

Through enhancements in temperature control, vibration isolation, error correction, and automated adjustment capabilities, our Ising machine, QBoson-CPQC-3Gen, demonstrates significant improvements in both stability and solution quality. Figure 1a illustrates the experimental setup of QBoson-CPQC-3Gen, including its measurement and feedback systems; additional details are provided in the Methods section. These improvements involve multidisciplinary optimizations in optics, structural design, and FPGA-based measurement-feedback control. A key upgrade is the refined temperature control system, which maintains optical fiber temperature within a narrow range over prolonged periods through a multi-layer enclosure design (Extended Data Fig. 1a). This enhancement allows the hardware to operate stably for extended durations without manual intervention, forming a critical foundation for applying the physical machine to CADD and other computationally intensive tasks. Furthermore, the machine supports 2000 qubits, sufficient for addressing approximately 70% of typical COPs^26^ (Extended Data Fig. 1b).

**Fig. 1.**
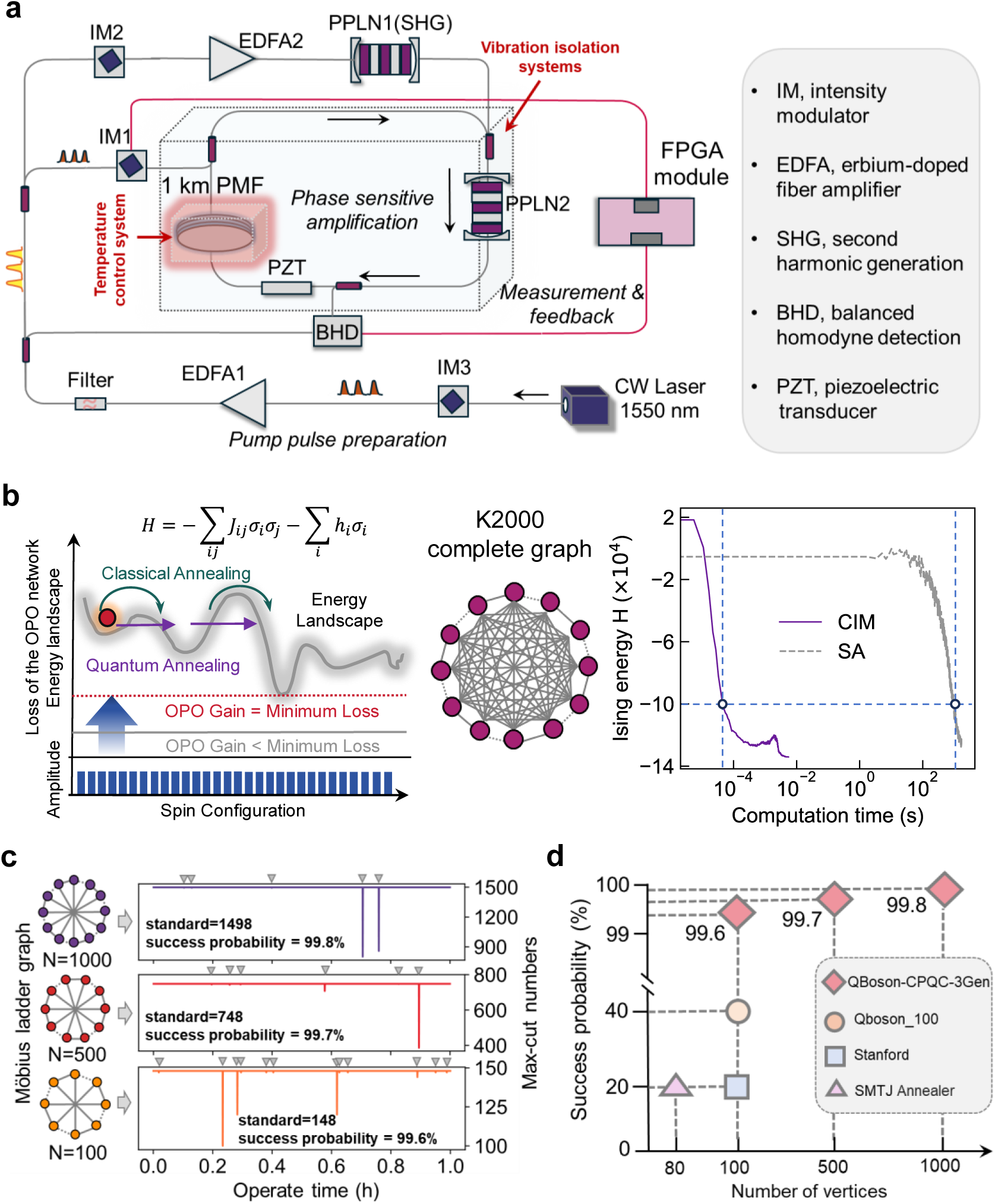
A stable 2000-node CIM. **a**. Setup of a coherent Ising machine (CIM) with measurement and feedback (MFB). **b**. The evolution schematic diagram of quantum annealing and simulated annealing, and the comparison of Hamiltonian evolution in solving the max-cut problem at 2000-node complete graph. **c**. Continuously solving Max-Cut problems of Möbius ladder graphs with 1000, 500, 100 nodes by QBoson-CPQC-3Gen in 1h. The grey triangles indicate the max-cut numbers did not meet the standard. **d**. The success probabilities of different quantum annealer in solving problems of different scales.

We first evaluated the computational performance of QBoson-CPQC-3Gen by solving maximum-cut (MAX-CUT) problems (28), which involve partitioning a graph into two subgraphs while minimizing the number of retained edges. We compared QBoson-CPQC-3Gen against simulated annealing (SA) on the MAX-CUT problem for a fully connected graph, K2000, containing 1,999,000 undirected edges (Fig. 1b). The evolution of Ising energy for both CIM and SA (Fig. 1b) highlights the substantial advantage of our machine in tackling large-scale combinatorial optimization problems. We further assessed stability by continuously solving MAX-CUT on Möbius ladder graphs^27^ (MLGs) of sizes 100, 500, and 1000 nodes—a standard benchmark for COP-solving machines. Although MAX-CUT on general graphs is NP-hard, MLGs admit deducible optimal solutions, providing a reference for evaluating machine performance^18, 19^. As shown in Fig. 1c, QBoson-CPQC-3Gen consistently produced solutions aligned with reference values across all graph sizes, achieving a single-shot success probability of no less than 99.5% for over one hour. Unlike prior evaluations that focused primarily on DOPO stability, our metric directly reflects the stability of the entire computational framework. Compared to several published physical machines^20, 28^, QBoson-CPQC-3Gen solves larger problem instances while sustaining high computational accuracy over longer periods (Fig.1d). In addition, compared to simulated annealing^9^ (SA) (Fig. 1b), a heuristic algorithm on classical computers imitating ground-state evolvement, not only does QBoson-CPQC-3Gen finds a better solution, but also saves the time for about 5 magnitudes. These results have marked an unprecedented milestone in CIM, and further ensured the potential application of QBoson-CPQC-3Gen in drug discovery scenarios.

Next, we applied QBoson-CPQC-3Gen to allosteric site prediction, protein-peptide docking, and molecular similarity calculation (Fig. 2). Building on prior success in solving graph COPs with CIMs^29^, we developed methods to transform CADD-specific COPs into graph representations compatible with QBoson-CPQC-3Gen, enabling a unified solution pipeline.

**Fig. 2.**
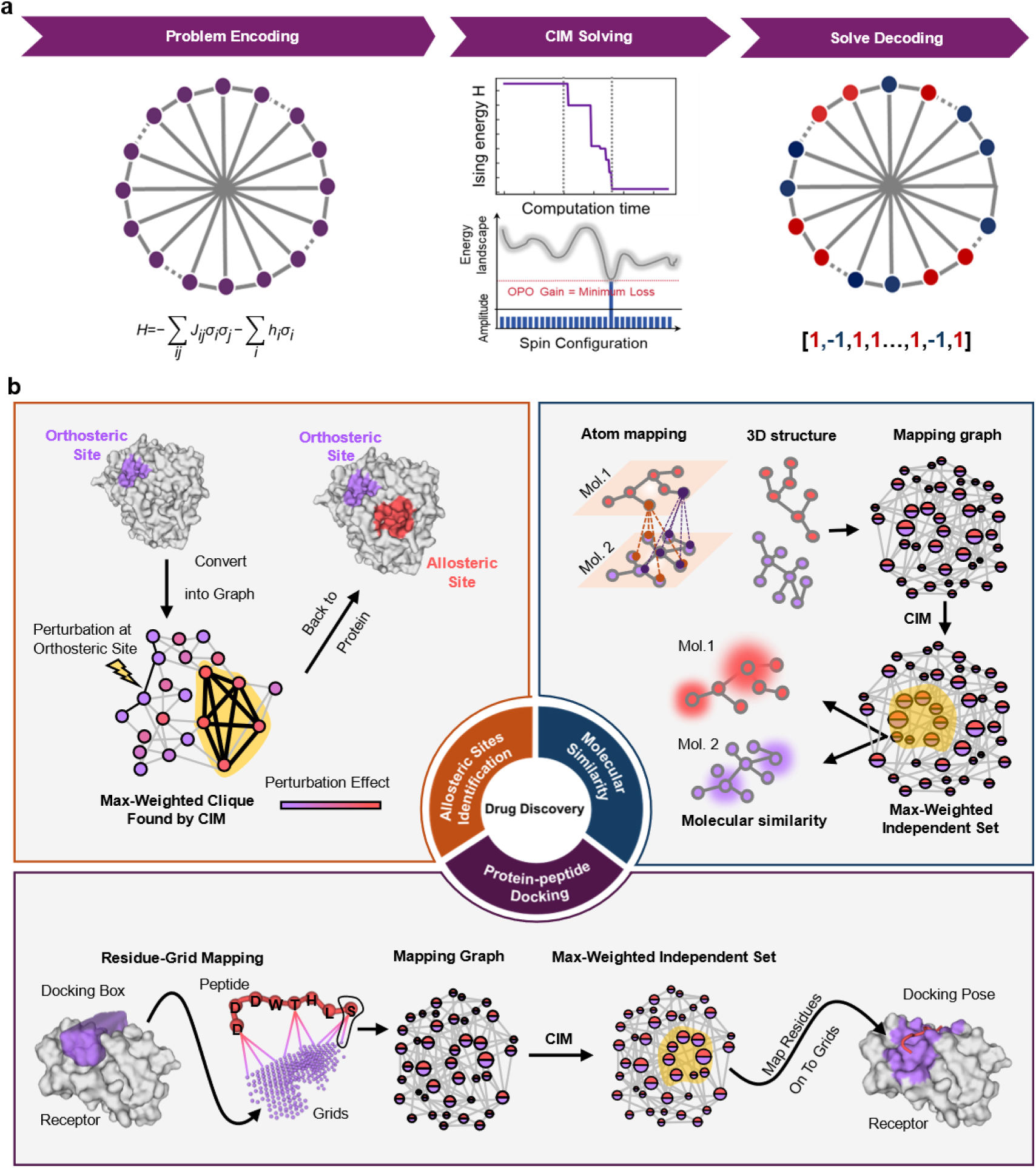
Schematic of quantum computing framework and three core drug discovery algorithms. **a.** A standard CIM-based programming solution process. **b.** Encoding details of three important quantum algorithms in drug discovery, including allosteric sites identification, protein-peptide docking, molecular similarity calculation.

### Allosteric Site Prediction by QBoson-CPQC-3Gen

Allosteric sites^30^ are druggable pockets on protein surface distinct from orthosteric (functional) sites (Fig. 3a). Drugs targeting allosteric sites are praised for higher selectivity and reduced toxicity^31–33^. However, identifying these sites remains a major challenge. Wet-lab discovery of allosteric sites^34–36^ are usually serendipitous and thus inefficient. Therefore, great expectations were given to computational methods. In a mathematical view, allosteric sites prediction is to find an optimal and spatially-continuous subset of protein residues. Previous computational methods typically rely on geometry-inspired local-search strategies, which first detect putative pocket regions and then evaluate whether each of them is allosteric^37–40^. While computationally efficient, t these approaches are constrained by the accuracy of the initial pocket detection^41^ (Extended Data Fig. 2a).

**Fig. 3.**
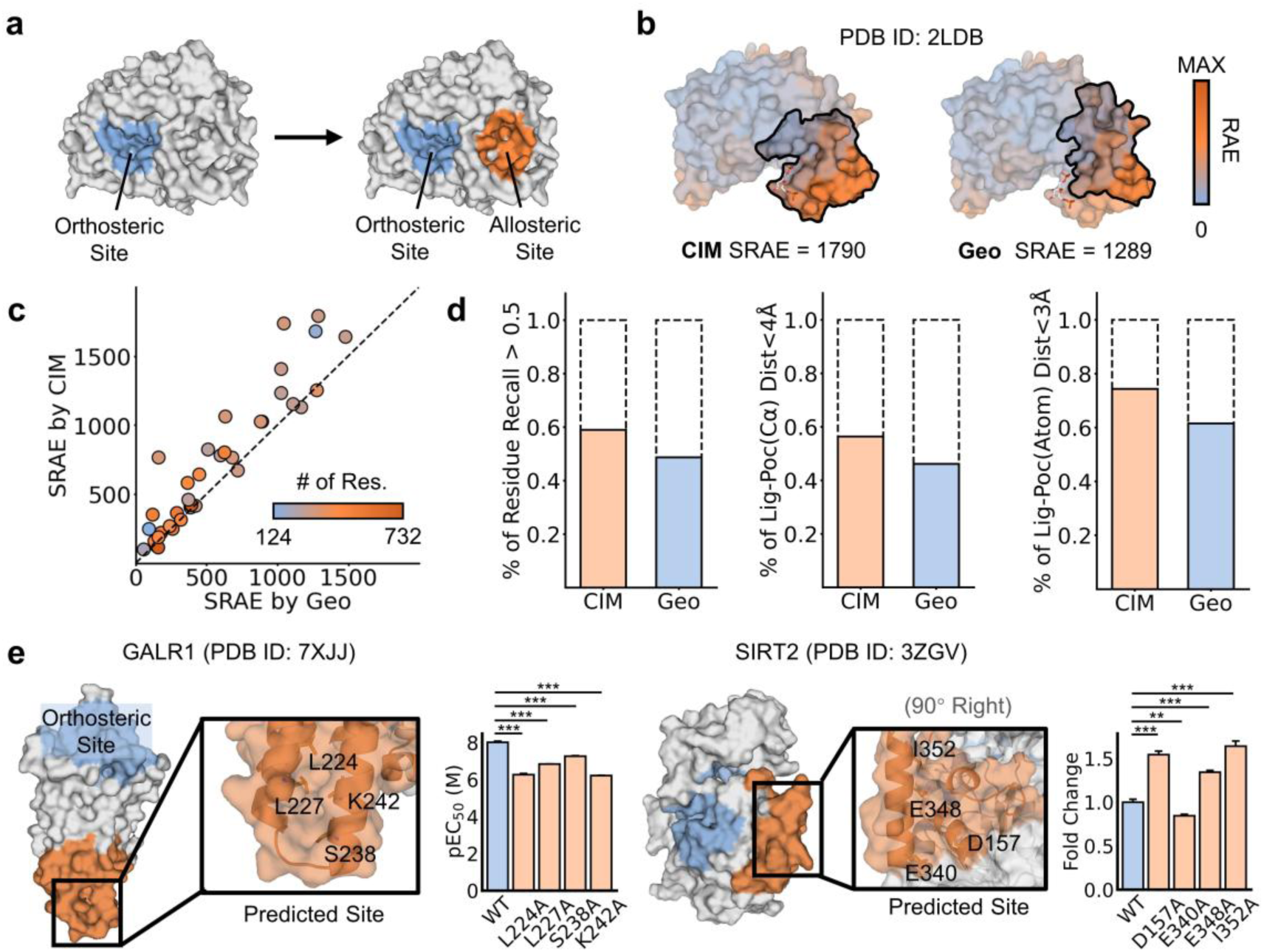
Allosteric site prediction with CIM. **a.** Illustration of allosteric site prediction. **b.** An example shows the failure of the geometric method (Geo) from like coherent Ising machine (CIM) because it doesn’t find a subgraph with lower sum of reversed allosteric effect (SRAE). Allosteric ligand is drawn in white sticks. **c.** Comparison of the SRAEs of subgraphs found by CIM and Geo. The dots are colored by the number of residues in protein. **d.** Ratio of allosteric sites predicted by CIM compared to Geo. e. Novel allosteric sites predicted by CIM (top row) and validations by site-directed mutagenesis (bottom row). Error bars means standard deviation. Statistics analysis are analyzed using Student’s t-test (normal and homo-variance) or Mann-Whitney test (otherwise). N=3. *: p<0.05. **: p<0.01. ***: p<0.001.

In this study, leveraging the high computational efficiency of QBoson-CPQC-3Gen, we implemented a global-search strategy for allosteric site prediction. Briefly, the protein structure is represented as a graph wherein residues serve as nodes, and edges connect residues within a specific distance. Node weights are assigned based on reversed allosteric effects (RAE), with higher values indicating a greater likelihood that the residue belongs to an allosteric site^38^. Allosteric site detection is then reformulated as finding the max-weighted clique (MWC), namely, identifying the fully-connected subgraph with the largest sum of RAE (SRAE). This graph problem could efficiently be solved by QBoson-CPQC-3Gen (Fig. 2).

We have benchmarked the algorithm on 39 proteins (Table S1, Extended Data Fig. 2b, Data S1) and taking geometry-inspired local-search (Geo)^42^ as comparison. Overall, QBoson-CPQC-3Gen achieved better performance in solving the MWC problem, identifying higher-weighted subgraphs in 79.5% of cases (Fig. 3c, Extended Data Fig. 2c). Furthermore, the CIM-based approach identified more known allosteric sites across both residue recall and ligand-distance metrics (Fig. 3d). A representative example (Fig. 3b) illustrates a case where Geo failed to predict the allosteric site due to its suboptimal performance in identifying the MWC (Fig. 3c).

We further applied QBoson-CPQC-3Gen to predict novel allosteric sites on two proteins including Galanin receptor 1 (GALR1) and Sirtuins 2 (SIRT2), which are important targets for depression^43^ and aging^44^, respectively. The predictions are shown in Fig. 3e. To validate these predictions, we performed alanine mutagenesis on key residues within each predicted site. Significant functional changes were observed in both cases, supporting the potential allosteric functionality of these regions (Fig. 3e, Table S2-S3, Extended Data Fig. 2d). Collectively, these results demonstrate that the global-search capability of QBoson-CPQC-3Gen significantly enhances performance in allosteric site detection.

### Peptide Docking by QBoson-CPQC-3Gen

Molecular docking^45^, a crucial technique in virtual drug screening, aimed at predicting the optimal binding pose of a ligand to its receptor. The process can be divided into two main steps: pose sampling and pose ranking^46^ (Fig. 4a). pose sampling constitutes a challenging combinatorial optimization problem (COP). While we previously developed a CIM-based approach for this step^23^, it relied on fixed ligand conformations, making it unsuitable for docking flexible molecules such as peptides^47–50^. In the current work, we have developed a flexible peptide docking method capable of handling conformational flexibility using QBoson-CPQC-3Gen.

**Fig. 4.**
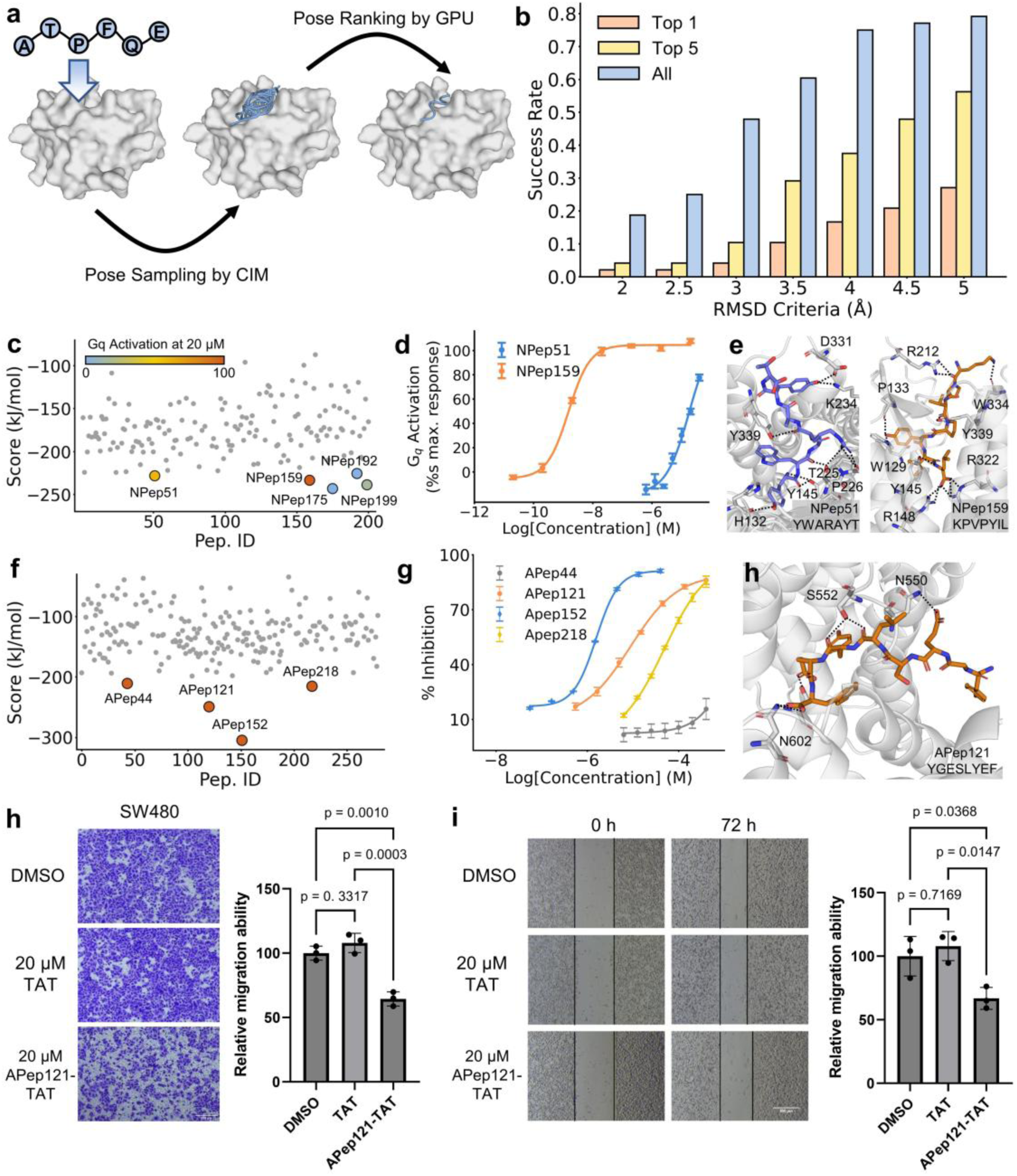
Peptide docking with CIM. **a.** Illustration of peptide docking. **b.** Ratio of successful docking with different criterial of root mean square distances (RMSDs) and number of poses selected. **c.** Result of virtue screening NTSR1 activators by CIM. **d.** Dose-depend response curve of the selected peptides from **c. e.** Docking pose of selected NTSR1 activators. **f.** Result of virtue screening APC-Asef inhibitors by CIM. **g.** Dose-depend response curve of the selected peptides from **f. h.** Docking pose of selected APC-Asef inhibitors. **i.** Transwell migration assays (SW480) of APep121-TAT, where TAT is the cell-penetrating peptide. **j.** Wound healing assays (SW480) of APep121-TAT. N=3 for all experiments. The error bars show the standard deviation. Significance tests were done using one-way ANOVA with normality and homo-variance checked.

Building on previous work^23^, the binding site is discretized into a grid, and pose sampling is formulated as the problem of assigning peptide residues to suitable grid points. To identify optimal placements, a fully connected graph is constructed where each node represents a possible residue–grid assignment, weighted by its corresponding binding energy^51^ . Edge weights encode inter-residue interaction energies and penalty terms. Penalties are applied if two nodes share the same grid point or residue, or if they lead to unrealistic peptide conformations. Thus, sampling low-energy binding poses for an N-residue peptide translates into searching for N-node subgraphs with minimal total weight, a task efficiently handled by QBoson-CPQC-3Gen (Fig. 2).

We evaluated this method on a dataset of 48 peptides comprising 5 to 9 residues (Table S4, Data S2) ^47, 52^. QBoson-CPQC-3Gen demonstrated strong performance in identifying low-energy binding poses (Extended Data Fig. 3a). Near-native poses (RMSD < 3 Å) were recovered in approximately 50% of cases, and in nearly 80% of cases a pose with RMSD < 5 Å was identified (Fig. 4b). Notably, low-RMSD poses were not exclusively enriched among the lowest-energy solutions, which may be attributed to the simplified residue-based energy function used. Overall, these results affirm the capability of QBoson-CPQC-3Gen in peptide pose sampling.

We next applied this approach to screen for peptide activators of neurotensin receptor type 1^53^, a target relevant to treating addictive behaviors. Following CIM-based sampling, poses were re-scored using an atom-level scoring function^54^ on a graph processing unit (GPU). The docking box was centered on the orthosteric site of PDB entry 8JPF^55^. The virtual screening was performed on a library of 200 hexapeptides (Fig. 4c, Table S5), with top five hits selected for experimental validations (Table S6–S8, Fig. S1–S10). Among these, NPep51 (YWARAYT) and NPep159 (KPVPYIL) activated both Gq-mediated signaling (Fig. 4d–e) and β-arrestin2 recruitment (Extended Data Fig. 3b), which are two downstream pathways of NTSR1. Notably, NPep159 exhibited nanomolar activity, highlighting its promise for further therapeutic development.

We also employed the combined CIM-GPU screening pipeline to identify peptide inhibitors targeting the protein–protein interaction (PPI) between adenomatous polyposis coli (APC) and Rho guanine nucleotide exchange factor 4 (Asef)^56^ (PPI), which is a promising strategy to inhibit colorectal cancer (CRC) migration. The screening was performed on a 5-8 peptide library (Fig. 4f, Table S9). The docking box is defined from the peptide in PDB 5Z8H. Top-4 candidates were selected, and APep121 (YGESLYEF) and APep152 SCESLYEF) were found with better EC_50_ values of 8.74 μM and 1.56 μM, respectively (Fig.4g-h, Table S10, Extended Data Fig. 3c, Fig. S11-S18). To assess their capacity to inhibit cancer migration in-cell, we conjugated the C-termini of the peptides to a cell-penetrating peptide (TAT). While APep152 lost activity upon conjugation, APep121-TAT significantly inhibited migration in CRC cell lines (SW480 and DLD-1) at 20 μM in both transwell migration assays (Fig. 4h, Extended Data Fig. 3d) and wound-healing assays (Fig. 4i, Extended Data Fig. 3e). These results demonstrate the successful application of QBoson-CPQC-3Gen in discovering peptides with in-cell efficacy.

### Similarity-Based Drug Screening by QBoson-CPQC-3Gen

Evaluating intermolecular similarity^57–59^ is another essential technique in virtue drug screening, particularly for identifying “me-better” compounds. One approach to calculate similarity involves mapping atoms from one molecule to another, with similarity defined as the average ratio of mapped atoms^60^ (Fig. 5a). This atom-mapping problem can be formulated as a combinatorial optimization problem (COP). To solve it, we represent the task as a graph where each node corresponds to a potential atom-atom match, weighted by its chemical compatibility. Edges are introduced between conflicting nodes, such as those sharing the same atom or violating geometric constraints, in order to prevent invalid mappings. The optimal mapping is then equivalent to finding the maximum-weighted independent set (MWIS), defined as the node set with the highest total weight and no connecting edges, a problem well-suited for coherent Ising machines (CIMs).

**Fig. 5.**
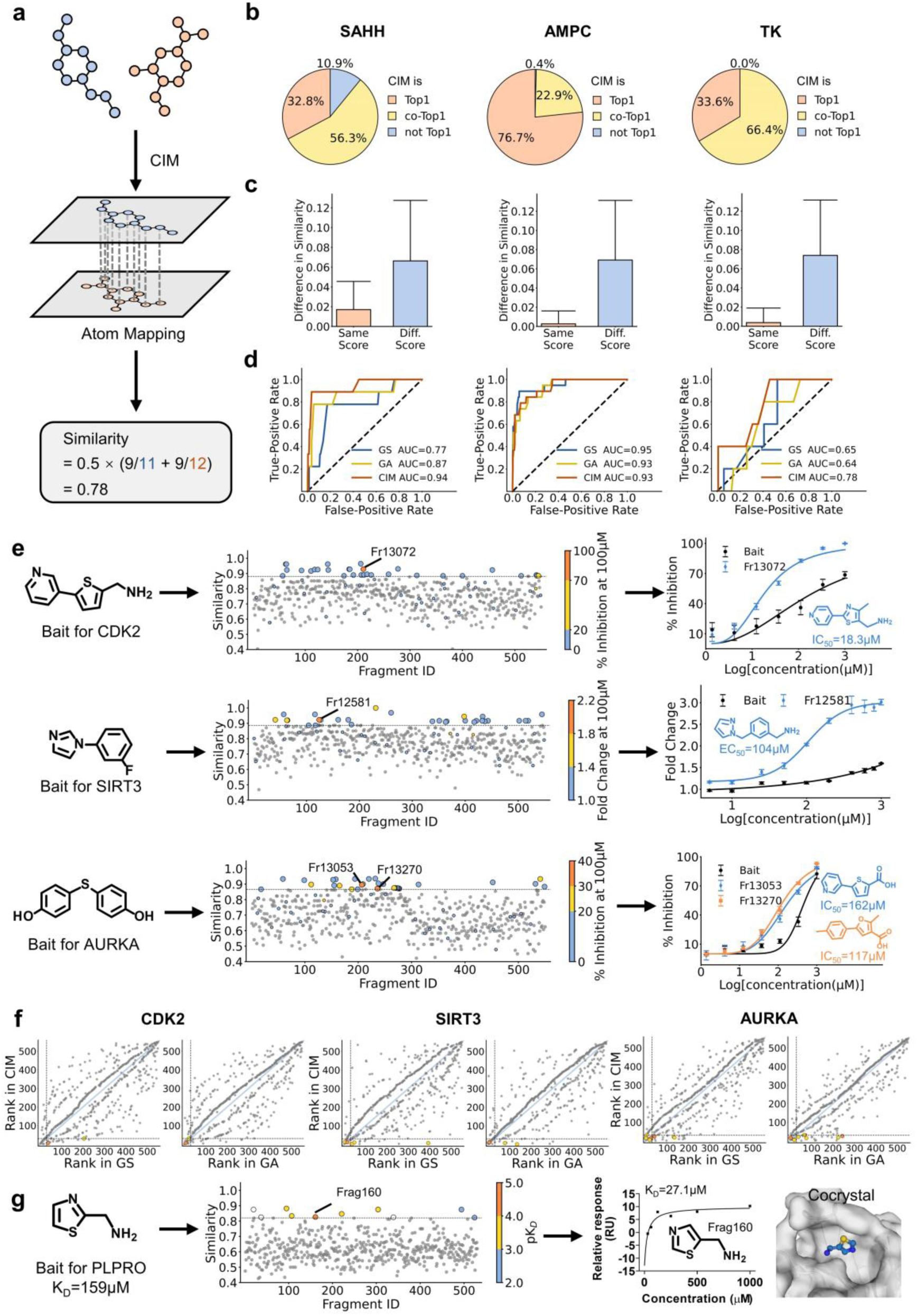
Molecular similarity calculation with CIM. **a.** Illustration of intermolecular similarity calculation. **b-d.** Benchmarking CIM-based similarity calculation on DUD dataset. Heuristic algorithms including greedy search (GS) and genetic algorithm (GA) are used as reference. **b.** Score of solved max-weighted independent set (MWIS). **c.** Influence of calculated similarity by MWIS score. **d.** Receiver operating curve (ROC) by the calculated similarity in classifications of molecules in DUD. **e** Ligand-based drug screening by CIM. Left: Workflow. Middle: Result. Top 30 fragments and a random selection of the rest molecules are tested by wet-lab experiments (shown by colors). Right: Dose-depend responses. **f.** Comparison of ranking difference when calculating similarity using CIM, GS and GA. Positive molecules in **e** are marked here with the same color. g. Screening binders of PLPRO using CIM. The best molecule is validated by SPR and crystal structures. N=3 for all experiments. The error bars show the standard deviation.

We benchmarked QBoson-CPQC-3Gen on 4 cases from the DUD dataset^60^, and its performance was compared to CPU-based heuristic algorithms, including greedy search^61^ (GS) (Fig. S19) and genetic algorithm^62^ (GA) (Fig. S20). In each case, similarity was computed between a query compound (active) and a set of both active and decoy molecules. Ideally, similarity scores should be higher for query-active pairs than query-decoy pairs. Starting from the MWIS solution, CIM matched or outperformed classical methods in 89.1% to 100% of pairs and achieved a better MWIS in 32.8% to 88.9% of cases (Fig. 5b, Extended Data Fig. 4b). Moreover, CIM reduced computation time by one to five orders of magnitude compared to CPU-based algorithms (Extended Data Fig. 4a). Notably, the runtime of GA and GS was highly sensitive to graph size, while CIM consistently completed within 0–4 ms for nearly all pairs, indicating its strong potential for large-molecule comparison. Slight variations in CIM solving time are likely attributable to random noise (Extended Data Fig. 1c). The MWIS solutions were then converted into similarity scores. We observed that different MWIS solutions with identical total weights could yield slightly different similarity values, and these variations were substantially smaller than those resulting from different MWIS weights (Fig. 5c), underscoring the importance of solution quality in similarity assessment. Finally, we used the derived similarity scores to classify query-active versus query-decoy pairs. For SAHH and TK targets, CIM achieved area under receiver-operating curve (AUC) values approximately 0.1 higher than GA or GS. For AMPC and HSP90, CIM performed comparably to classical methods (AUC difference < 0.02), likely due to error cancellation (Fig. 5d). Overall, these results demonstrate that CIM outperforms classical heuristic algorithms in molecular similarity calculation.

We further investigated whether QBoson-CPQC-3Gen could facilitate real-world “me-better” compound discovery. Using previously identified dose-dependent fragments targeting Aurora A kinase^63^ (AURKA), Sirtuin 3^64^ (SIRT3) and cyclin-dependent kinase^65^ (CDK2), which are famous targets in treatments of breast cancer, cardiovascular diseases and leukemia, respectively. These fragments were adopted as baits for searching structure-similar molecules in a fragment database using QBoson-CPQC-3Gen (Fig. 5e). The top 30 most similar fragments were purchased and tested experimentally, alongside 30 randomly selected fragments from the rest as a negative control. As shown in Fig. 5f, 2 to 8 moderately (yellow) or highly (orange) active fragments were identified among the top CIM-derived hits, whereas the control group contained only 1-2 moderately active and no highly active fragments. Highly active compounds shared strong similarity (∼0.9) with the bait molecules, though scaffold modifications such as atom insertions or substitutions were observed. Dose-response measurements (EC₅₀ or IC₅₀) of these newly identified hits were comparable to or better than those of the original bait fragments (Tables S11–S13). For comparison, we repeated the screening using GS and GA. The similarity rankings produced by these methods diverged from CIM’s results (Fig. 5f), and many active fragments identified by CIM were ranked beyond the top 30 by GS or GA (dashed lines), indicating that conventional methods would have missed these hits. Further analysis suggested that this discrepancy stems from the suboptimal MWIS solutions found by GA and GS, which led to inaccurate similarity scores (Extended Data Fig. 5b–c).

We performed the same screening workflow on Papain-like protease^66^ (PLPRO), a replicase in SARS-Cov-2, with a bait molecule (Extended Data Fig. 5d) to query a new fragment library (Fig.5g). Top 10 most similar molecules were selected for experimental validation via surface plasmon resonance (SPR). Five compounds exhibited micromolar dissociation constants (K_D_), with Frag160 showing a K_D_ of 27.1 μM (Fig. 5g, Extended Data Fig. 5e, Fig. S21, Table S14). Co-crystal structures confirmed that Frag160 binds to the same allosteric site as GLR0617 (Fig. 5g, Extended Data Fig. 5g, Table S16), a known PLPRO inhibitor^66^. In summary, we have demonstrated that CIM-based approaches offer enhanced performance in molecular similarity evaluation and can be effectively applied in practical ligand-based drug screening campaigns.

## Discussion

A major challenge in computer-aided drug discovery (CADD) lies in the repeated solving of large-scale combinatorial optimization problems (COPs)^4^. Conventional approaches such as heuristic algorithms^9^ are often time-consuming and yield suboptimal solutions. Cutting-edge machines like coherent Ising machines^18^ (CIMs) offer the potential to solve large-scale COPs rapidly and accurately. However, limitations in operational stability have hindered their adoption in CADD. Furthermore, a lack of efficient methods for encoding CADD-related COPs into CIM-compatible formats has posed an additional barrier.

To address these challenges, we developed QBoson-CPQC-3Gen, a new-generation CIM designed with enhanced stability through improved fiber vibration isolation and precision temperature control. This machine can COPs involving up to 2000 variables while continuously outputting high-quality solutions for over an hour. Its exceptional stability in handling large-scale COPs underscores its potential not only for CADD but also for other computationally intensive applications. On the software front, we introduced a graph-based encoding strategy to translate CADD-specific COPs into CIM-readable formulations. Graphs align naturally with the structure of CIMs, where nodes correspond to qubits and edges represent inter-qubit couplings. This structural similarity builds upon prior work in solving subgraph searching problems with CIMs^29^. Therefore, it is intuitive to convert COPs in CADD into well-studied graph problems. Using this approach, we realized solution of three key tasks in CADD on CIM, including allosteric site detection, protein-peptide docking, and molecular similarity calculation. Central to our encoding strategy was the representation of fundamental problem elements, such as protein residues or residue-grid assignments, as nodes within a graph model. The CIM-based solutions achieved speedups of several orders of magnitude while delivering superior solution quality compared to classical heuristic algorithms, demonstrating a tangible computational advantage often associated with quantum-inspired systems.

More importantly, we present pioneering examples of applying quantum computing hardware to real-world drug discovery, leading to the identification of two previously unreported allosteric sites, two novel peptide modulators, and five small-molecule modulators. These findings were rigorously validated through *in vitro* assays, in-cell experiments, and crystallographic analyses.

In conclusion, through concurrent advancements in both hardware and encoding methodologies, we have established a pipeline for solving COPs arising at multiple stages of drug discovery. We recognize that the current scale of available spins and computational precision may constrain applications to more complex scenarios, particularly in modeling intermolecular relationships, where the required number of qubits scales quadratically with system size. Therefore, future efforts should focus on further increasing the spin capacity and operational precision of CIMs to broaden their applicability in computational biology and drug design.

## Methods

### Setup of coherent Ising machine

A continuous-wave laser with a central wavelength of 1550.1 nm was modulated by an intensity modulator (IM3) to generate a pulse train at a repetition rate of 1 GHz and a pulse width of approximately 30 ps. The pulse train was amplified using an erbium-doped fiber amplifier (EDFA1) and split into two paths. One path served as the local oscillator input for a balanced homodyne detector (BHD). The other path was further divided: one portion passed through IM1 to carry feedback signals from the FPGA, serving as injection light; the other portion was fed into IM2 to control the pump amplitude. These pulses were then amplified by EDFA2 and directed into a periodically poled lithium niobate (PPLN) waveguide module. Second-harmonic generation (SHG) in PPLN1 produced a 775 nm pulse train, which was injected into an optical cavity containing PPLN2 (functioning as a phase-sensitive amplifier, PSA), two optical couplers, 1 km of optical fiber, and a piezoelectric transducer (PZT) for cavity stabilization.

### Temperature control and vibration isolation system of optical path

To maintain thermal stability, the 1 km fiber was housed in a rubber-plastic wrapped storage barrel filled with water—a high heat-capacity medium—to buffer against temperature fluctuations. For vibration isolation, the free-space optical path was enclosed using thermal insulation and damping materials to minimize environmental mechanical and thermal disturbances.

### Solving combination optimization problems with CIM

We outline a general workflow for solving combinatorial optimization problems (COPs) using a coherent Ising machine (CIM). Mathematical details for each specific CADD problem are provided in the Supplementary Text under “Details of the problem encoding schemes.” First, the COP is transformed into a subgraph search problem by representing basic problem elements as nodes: residues for allosteric site detection; residue–grid assignments for protein-peptide docking; and atom-atom correspondences for molecular similarity evaluation. A quadratic unconstrained binary optimization (QUBO)^67^ model is then established, where each node is associated with a binary variable *x*_*i*_ ∈ {0,1}, indicating whether the node is included in the solution subgraph. The graph problem is thus reformulated as minimizing a quadratic function. To interface with the CIM, the QUBO model is converted into an Ising model, which is also a quadratic binary minimization model, but with spin variables *s*_*i*_ ∈ {−1,1}. The Ising model is input into QBoson-CPQC-3Gen, with couplings configured to represent the problem’s interaction terms. The system evolves to its ground state, and the resulting spin configuration is read out to recover the solution to the Ising model, which corresponds to the optimal subgraph and thereby the solution to the original COP.

### Reversed allosteric effect

The reversed allosteric effect (RAE) was computed as previously described^38^to estimate the likelihood of a residue being part of an allosteric site. Briefly, the protein bound to an orthosteric ligand was coarse-grained into two models: an *apo* model with only Cα atoms, and a *holo* model that also included heavy atoms of the ligand. Ensembles for both models were generated via normal mode analysis^68^. Potential energies were computed for each frame and projected per residue. The RAE for a residue was defined as the mean absolute deviation of its energy between the *apo* and *holo* ensembles.

### Predict allosteric sites with a geometric approach

Putative pockets were identified using Fpocket^42^. To ensure comparability, each pocket was adjusted such that no two residues exceeded 28 Å apart (Extended Data Fig. 2b). The sum of RAE (SRAE) values was computed for each pocket, excluding residues within 12 Å of any orthosteric ligand atom. The pocket with the highest SRAE was predicted to be the allosteric site.

### Benchmarking of allosteric site prediction

The AlloReverse^38^ training and test sets were filtered to include only PDB entries containing both orthosteric and allosteric ligands within a single chain. Allosteric sites were defined as residues within 9 Å of the allosteric ligand. Entries where any allosteric site Cα atom was within 12 Å of the orthosteric ligand were excluded, resulting in a final set of 39 proteins. Predictions were evaluated based on residue recall rates and distances between predicted sites and the allosteric ligand.

### Protein expression of GALR1 and bioluminescence resonance energy transfer (BRET) assay for G protein dissociation

The BRET probe plasmids (Gαq-RLuc8, Gβ3, Gγ9-EGFP) utilized in this study were from the TRUPATH kit, which was provided by Bryan Roth laboratory (Addgenekit#1000000163). Human *GALR1* was cloned into a modified pcDNA3.1 (+) vector with a HiBiT tag (Promega) at the N-terminus. GALR1 mutations were introduced via site-directed mutagenesis using the Mut Express II Fast Mutagenesis Kit V2 (Vazyme). HEK293T cells (ATCC; routinely tested for mycoplasma) were transiently co-transfected with either wild-type or mutated GALR1 receptors alongside G protein BRET probes using the ExFect Transfection Reagent (Vazyme) according to the manufacturer’s protocol. After 24 h, the cells were reseeded into white opaque-bottom 96-well plates at a density of 20,000 cells/100 μL per well, then cultured in DMEM with 10% FBS at 37 °C and 5% CO₂ for another 16 h. Subsequently, the culture medium was replaced with Hank’s Balanced Salt Solution, followed by stimulation with Galanin^1–16^ at varying concentrations for 5 min. The BRET fluorescence signal was then detected after addition of 5 μM Coelenterazine 400a (Maokangbio) using a Synergy Neo plate reader (BioTek, USA), with emission filters at 410 nm and 515 nm.

### Protein expression and purification of SIRT2

The N-terminally His-tagged wild-type (WT) SIRT2 protein was expressed in Escherichia coli Rosetta (DE3) cells. Site-directed mutagenesis (Y150A, R153A, L154A, D157A, E340A, L344A, E348A, I352A) was performed using the Mut Express II Fast Mutagenesis Kit V2 (Vazyme) according to the manufacturer’s protocol. Transformed colonies were selected on LB agar plates supplemented with kanamycin (50 μg/mL) and chloramphenicol (34 μg/mL). A single colony was inoculated into 20 mL of LB medium containing the same antibiotics and grown overnight at 37°C with shaking. The culture was then expanded to 1L in fresh LB medium and incubated until the optical density at 600 nm (OD600) reached 0.6-0.8. Protein expression was induced by adding 0.2mM isopropyl β-D-1-thiogalactopyranoside (IPTG), followed by incubation at 12°C for 16-18 h. Cells were harvested by centrifugation and resuspended in lysis buffer (50 mM Tris-HCl, pH 8.0, 500 mM NaCl, 5% glycerol, 1 mM phenylmethylsulfonyl fluoride (PMSF), 1 mM dithiothreitol (DTT), and 1× protease inhibitor cocktail). The clarified lysate was purified by Ni-NTA affinity chromatography (HisTrap FF), followed by size-exclusion chromatography (Superdex S200 Increase 10/300 GL) in buffer (50 mM Tris-HCl (pH 8.0), 137mM NaCl, 2.7mM KCl, 1mM MgCl2). Protein purity was verified by SDS-PAGE before concentration.

### Fluorescence detection (FD) assay for SIRT2

For the SIRT2 basal enzymatic activity, 200 nM SIRT2 was incubated with 80 μM substrate peptide (RHKK(ac)-AMC) and 500μM NAD+ at a final DMSO concentration of 1% in assay buffer (50 mM Tris-HCl, pH 8.0, 137 mM NaCl, 2.7 mM KCl, 1 mM MgCl2). The reactions were carried out at 37°C for 45min, terminated with 6 mM nicotinamide, and developed with 5.5 mg/mL trypsin for 30min at rm. The fluorescence intensity was measured with a microplate reader ((SynergyNeo, BioTek) at excitation and emission wavelengths of 360 nm and 460 nm, respectively.

### Benchmarking the sampling power in peptide docking

The test set of HPepDock^47^ and another benchmarking set^52^ are collected. To meet the requirements in problem scale, only cases with 5 to 9 residues are chosen, and cases whose precision of ising matrix above 14 were removed, yielding a dataset of 48 complexes. Redocking was performed with 1000 CIM samples per case. Performance was evaluated by the RMSD between docked and crystal poses.

### Atom-level scoring function

All-atom models were reconstructed from coarse-grained poses using cg2all^69^. Clashes could exist in all-atom models, so that a minimization followed by a 10-ps equilibration simulation is performed by OPENMM^70^ with FF14SB force^54^ and GBN2^71^ implicit solvent model. Binding energies were computed via MM/GBSA^72^ over a subsequent 10 ps simulation and averaged for scoring.

### Protein expression of NTSR1 and Bioluminescence resonance energy transfer assay

HEK293T cells (ATCC; the cells were routinely tested for mycoplasma contamination) were used for all experiments. The cells were cultured in Dulbecco’s modified Eagle’s medium (DMEM) supplemented with 10% FBS. In brief, cells were seeded at a density of 3–4 million cells per dish in 10-cm dishes. After 18-24 h, cells were transfected at a confluency of 70–80%, using a 1:1:1:1 DNA ratio of NTSR1: Gαq–RLuc8: Gβ: Gγ–GFP2 (2 μg per construct for 10-cm dishes, the plasmids encoding subunits of the heterotrimeric G protein complex named TRUPATH was a gift from Bryan Roth, Addgene kit #1000000163), or using a 1:1:10 DNA ratio of NTSR1–RLuc8: GRK2: GFP2-β-arrestin2 (total 8 μg for 10-cm dishes). ExFect Transfection Reagent (Vazyme) was used to complex the DNA at a ratio of 2 μl of ExFect per μg of DNA, Opti-MEM (Gibco) at a concentration of 10 ng of DNA per μl of Opti-MEM. The next day, cells were reseeded in white opaque-bottom 96-well assay plates (Beyotime) at a density of 30,000–50,000 cells per well.

At 24 h later, culture medium was replaced with 80 μl of assay buffer (HBSS supplemented with 20 mM HEPES, pH 7.4) containing 5 μM coelenterazine 400a (Nanolight Technologies). After a 2-min equilibration period, cells were treated with 20 μl of peptides prepared in drug buffer at serial concentration gradient for an additional 5 min. Plates were then read in a Synergy Neo plate reader (BioTek) with 410-nm (RLuc8–coelenterazine 400a) and 515-nm (GFP2) emission filters, at integration times of 1 s per well. BRET2 ratios were computed as the ratio of the GFP2 emission to RLuc8 emission. Normalization was done using the reference ligand (NTS^8–13^ for NTSR1) as a divisor, which means BRET2 ratio at [reference ligand]0 was designated as 0%, BRET2 ratio at [reference ligand]max as 100%.

All peptides (synthesized by GenScript) were dissolved in DMSO to prepare 10 mM stock solutions. For the primary screening, each peptide was tested at a final concentration of 20 μM in both G-protein dissociation and β-arrestin recruitment assays to evaluate potential agonist and antagonist activities.

### Protein expression and purification of APC

APC (R303–D739) construct was cloned into pET28a vector with an N-terminal hexahistidine tag and expressed in E. coli strain BL21 (DE3). Cells were initially cultured in LB broth supplemented with kanamycin (50 μg/mL). After 6 h, cells were inoculated into large-scale ZYM-5052 medium supplemented with kanamycin (100 μg/mL) and incubated at 37 ℃ for 6 h, followed by incubation at 25 ℃ for 18 h with shaking at 250 rpm. Cells were harvested by centrifugation at 7000 × g for 15 min. Cell pellets were resuspended in lysis buffer (25 mM Tris-HCl, pH 8.0, 300 mM NaCl, and 20 mM imidazole) and lysed using a high-pressure homogenizer. Lysates were centrifuged at 23,000 rpm for 30 min, and loaded onto a HisTrap FF Ni-NTA column (Cytiva), washed with lysis buffer, and eluted with elution buffer (25 mM Tris-HCl, pH 8.0, 300 mM NaCl, and 500 mM imidazole). Samples were concentrated and exchange to storage buffer (50 mM HEPES, pH 7.5, 300 mM NaCl, 1 mM EDTA). The purified APC was concentrated, aliquoted and flash frozen for -80 ℃ storage.

### FP-based competitive assay for peptide inhibitors

The assays were run in triplicate on 96-well plates (Corning 3650). The final assay volume was 100 μL, containing 25 nM APC protein, 20 nM tracer (Cbz-AGESLYEK-FITC-NH2) and decreasing concentrations of peptides (7-point 2-fold serial dilution). All components were diluted in FP buffer (50 mM HEPES, pH 7.5, 300 mM NaCl, 1 mM EDTA, 1 mM DTT). The FP signals were measured at room temperature using a Synergy neo microplate reader (BioTek) with fluorescence excitation and emission wavelengths of 485 and 528 nm, respectively. Control wells containing APC and tracer (zero displacement), or free tracer only (maximum displacement) were also included. Data were normalized to control values to obtain percentage inhibition. The dose-response curves were fitted by nonlinear regression analysis using scipy. The IC_50_ values of peptides were calculated as the mean of three independent experiments.

### Wound healing assay

SW480 and DLD-1 cells were seeded in 6-well plates at densities of 1.25 × 106 cells/well and 1× 106 cells/well, respectively, and cultured to near confluence. Monolayers were scratched using a 200 μL sterile pipette tip, washed three times with PBS to remove detached cells, and incubated in DMEM supplemented with 2% FBS with or without test compounds (final 20 μM, 0.2% DMSO). Phase-contrast images were acquired at 0 h and indicated times using a Nikon Eclipse Ti2 inverted microscope, and wound closure was quantified by measuring relative wound area using ImageJ (NIH).

### Transwell migration assay

Cell migration assays were performed using 24-well plates with 8 μm pore size transwell inserts (Corning 3422). SW480 and DLD-1 cells were serum-starved for 8 h, and then digested with TrypLE (Invitrogen) and resuspended in serum-free medium. A total of 200 μL cell suspension (8× 104 cells) was added to the upper chamber, and 600 μL DMEM with 10% FBS was placed in the lower chamber. Compounds (final 20 μM, 0.2% DMSO) were added to both upper and lower chambers. After 48 h incubation, non-migrated cells were removed using cotton-tipped applicators, and migrated cells were washed twice with PBS, fixed with 4% paraformaldehyde for 20 min, and stained with crystal violet for 20 min. Inserts were washed, air-dried, and imaged using a Nikon Eclipse Ti2 inverted microscope. Migrated cells were quantified using ImageJ (NIH).

### Greedy search

We performed an ensembled version of greedy search^61^ to solve the maximum weighted independent set (MWIS) problem. In brief, three strategies of greedy search are applied to one graph, resulting in three independent sets. The set with the largest sum of weights is accepted as the final solve. As shown in fig. S5, the three strategies used include: (1) iteratively removing nodes with small weight and big degree until the graph is independent; (2) iteratively picking out nodes with big weight and small degree while deleting their conjoint nodes until the graph is empty; (3) similar to (2), but the weights of adjacent nodes are considered to adjust the degree value. The greedy search is written in Numba-accelerated Python codes^73^ and performed on one core of an AMD EPYC 7H12 CPU.

### Genetic algorithm

We performed an ensembled version of genetic algorithm^62^ to solve the MWIS problem. As illustrated in fig. S6, 100 chromosomes, each of which represents nodes selected, are randomly generated. Chromosomes representing invalid independent sets are fixed by removing nodes with smaller weights or larger degrees. Chromosomes with top 2 sum of weights are preserved, while the rest are randomly processed with crossover and mutation. Fixations, crossovers, and mutations are repeated for 100 rounds for each type of fixation strategy, and a best independent set is selected. The genetic algorithm is written in Numba-accelerated Python codes^73^ and performed on one core of an AMD EPYC 7H12 CPU.

### Benchmarking intermolecular similarity calculations

The DUD dataset^60^ is downloaded from https://dud.docking.org/jahn/. Four cases, including SAHH, AMPC, TK and HSP90 were selected. In each case, molecules with above 25% deviation in heavy atom number compared to the query molecule are removed because they do not fit the mapping-based similarity calculation. The performances are evaluated by the classification power between the positive and negative molecules in each case.

### Similarity-based drug screening

A dataset of 550 molecules were randomly selected from the High Solubility Fragment Library provided by TargetMol. Similarities were calculated between molecules in the libraries and bait molecules. Top 30 most similar molecules were further validated in wet lab while another 30 molecules were also tested as negative controls.

### AURKA Protein Expression and Purification

The gene encoding human AURKA (122-403) was subcloned into pET30b(+) vector with an N-terminal hexahistidine tag and expressed in *E. coli* BL21(DE3)^74^. Cells were initially cultured in LB broth supplemented with kanamycin (50 μg/mL). After 6 h, cells were inoculated into large-scale ZYM-5052 medium supplemented with kanamycin (100 μg/mL) and incubated at 37 ℃ for 6 h, followed by incubation at 25 ℃ for 18 h with shaking at 250 rpm. Cells were harvested by centrifugation at 7000 × g for 15 min. Cell pellets were resuspended in lysis buffer (50 mM Tris, pH 8.0, 500 mM NaCl, 10% glycerol, 20 mM imidazole, 1 mM TCEP) and lysed using a high-pressure homogenizer. Lysates were centrifuged at 23,000 rpm for 30 min, and loaded onto a HisTrap FF Ni-NTA column (Cytiva), washed with lysis buffer, and eluted with elution buffer (50 mM HEPES, pH 7.5, 200 mM NaCl, 10% glycerol, 500 mM imidazole, 1 mM TCEP). Samples were desalted into desalting buffer (20 mM HEPES, pH 7.5, 300 mM NaCl, 10% glycerol, 1 mM TCEP). Samples were diluted into buffer A (20 mM HEPES, pH 7.5, 100 mM NaCl, 10% glycerol, 1 mM TCEP), and further purified by cation exchange chromatography using a HiTrap SP column (Cytiva), and eluted with buffer B (20 mM HEPES, pH 7.5, 1 M NaCl, 10% glycerol, 1 mM TCEP). Samples were concentrated and further purified by size exclusion chromatography (SEC) using a Superdex 75 increase 10/300 GL column (Cytiva) in SEC buffer (50 mM Tris, pH 7.5, 150 mM NaCl, 5 mM MgCl_2_, 1 mM TCEP). The purified Aurora A was concentrated, aliquoted and flash frozen for -80 ℃ storage.

### SIRT3 protein expression and purification

SIRT3 with an N-terminal histidine tag, was expressed in Escherichia coli using a pET28a-based construct (SIRT3-118-399). This expression and purification process followed well-established protocols described in the literature^75^. Initially, expression in E. coli was induced by 0.5 mM isopropyl 1-thio-β-D-galactopyranoside (IPTG, I5502, Sigma, America), followed by a purification sequence that began with centrifugation and subsequent Ni-NTA bead clarification.

### CDK2 protein expression and purification

To generate CDK2 phosphorylated on T160 (p-CDK2), the gene encoding human CDK2 (1-298) was subcloned into pRSFDuet1 vector with an N-terminal hexahistidine tag and coexpressed with GST-tagged yeast CAK in E. coli BL21(DE3)^65, 76^. The culture and induction methods were the same as described for Aurora A. Cell pellets were resuspended in lysis buffer (50 mM Tris, pH 8.0, 500 mM NaCl, 10% glycerol, 1 mM TCEP) and lysed using a high-pressure homogenizer. Lysates were centrifuged at 23,000 rpm for 30 min, and loaded onto a HisTrap FF Ni-NTA column (Cytiva), washed with lysis buffer, and eluted with elution buffer (1 × PBS, pH 7.4, 500 mM NaCl, 10% glycerol, 500 mM imidazole, 1 mM TCEP). Samples were concentrated and further purified by size exclusion chromatography (SEC) using a Superdex 75 increase 10/300 GL column (Cytiva) in SEC buffer (50 mM HEPES, pH 7.5, 150 mM NaCl, 10 mM MgCl2, 1 mM EGTA, 1 mM TCEP). The purified p-CDK2 was concentrated, aliquoted and flash frozen for -80 ℃ storage.

### Kinase assays for AURKA and CDK2

Kinase activity was measured in ProxiPlate 384 shallow well plates (Revvity) using ADP-Glo Kinase Assay kit (Promega). Kinase reactions were performed in duplicate in assay buffer (20 mM HEPES pH 7.4, 20 mM NaCl, 1 mM EGTA, 10 mM MgCl_2_, 0.02% Tween-20, 0.1 mg/mL BSA, 50 μM DTT). In the primary screening, fragment compounds at 10 mM were diluted 20-fold in assay buffer to obtain a working solution at 5% DMSO. For AURKA, 1 μL of the working solution was pre-incubated with 2 μL of Aurora A (10 nM final) at room temperature for 30 min. Reactions were initiated by adding 2 μL of a mixture of TPX2 (residues 1-43; 10 nM final), kemptide (50 μM final) and ATP (50 μM final), and incubated at room temperature for 60 min. For CDK2,1 μL of the working solution was pre-incubated with 2 μL of a mixture of CDK2 (5 nM final) and peptide substrate HHASPRK (100 μM final) at room temperature for 30 min. Reactions were initiated by adding 2 μL of a mixture of cyclin A2 (5 nM final) and ATP (200 μM final), and incubated at room temperature for 30 min. Reactions were then quenched with ADP-Glo reagent according to the manufacturer’s instructions. Luminescence was measured using a Synergy neo plate reader with an integration time of 1 s, and normalized to no-inhibitor DMSO controls. Measurements of dose-dependent responses were performed on 3 distinct samples.

### Fluorescence detection (FD) assay for SIRT3

The SIRT3 activity assay was conducted according to methods previously described^64^. Briefly, 250 nM of SIRT3 was incubated with 10 μM of the substrate peptide (Z-(Ac) Lys-AMC) and 1 mM NAD^+^ (V900401, Sigma, America) either in the absence or presence of the tested compounds, with a final DMSO concentration of 1%. After incubating at 37°C for 30 minutes, the reaction was terminated using 10 μM Trypsin (T4799, Sigma, America) and 2 mM NAM (72340, Sigma, America). Fluorescence was then measured by exciting the fluorophore at 360 nm and detecting the emitted light at

460 nm using a fluorometric plate reader (SynergyNeo, BioTek, America). Measurements of dose-dependent responses were performed on 3 distinct samples.

### Estimation of EC_50_ or IC_50_

Mean values of response with their doses were used to fit the following equations by scipy. EC_50_ or IC_50_ were then extracted.

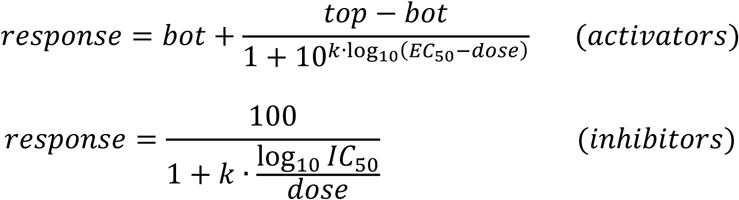

### Protein Expression and Purification of PLPRO

The coding sequences of SARS-CoV-2 PLPRO (residues 746–1060, wild-type and C111S mutant) were cloned into pET-28a and pET-21b vectors, respectively, and transformed into E. coli BL21 (DE3) cells. Bacterial cultures were grown at 37 °C until OD600 reached 0.6-0.8. Protein expression was induced with 0.5 mM IPTG, followed by overnight incubation at 18 °C. Cells were harvested by centrifugation and resuspended in lysis buffer (20 mM HEPES, 150 mM NaCl, 5 mM DTT, pH 7.5). Cells were lysed by sonication and the lysate was centrifuged at 13,000 rpm for 20 min at 4°C to remove cell debris. The protein was purified by Ni-NTA affinity chromatography. Further purification was performed by size-exclusion chromatography using a Superdex 200 column equilibrated with 20 mM HEPES (pH 7.5), 100–150 mM NaCl, and 5 mM DTT. Purified proteins were concentrated to 10–20 mg/mL, flash-frozen, and stored at –80 °C.

### Surface plasmon resonance

SPR experiments were performed on a Biacore 8K instrument (Cytiva). PLPRO with a N-terminal 6×His tag was immobilized on a CM5 sensor chip at a concentration of 50 μg/mL in 10 mM sodium acetate coupling buffer (pH 5.5 for PLpro and pH 5.0 for IDH1) to a surface density of 11,000 Response Units (RU) by amine coupling chemistry. The surface was first activated with a 1:1 mixture of EDC/NHS (contact time: 420 s, flow rate: 10 μL/min), followed by the immobilization of IDH1, then deactivated with 1 M ethanolamine (pH 8.5). Binding experiments were performed at 25 °C using a running buffer of 10 mM Na_2_HPO_4_, 1.8 mM KH_2_PO_4_, 137 mM NaCl, 2.7 mM KCl, 0.05% Tween 20, 2% DMSO, pH 7.4. Compounds were 2-fold serially diluted using the running buffer and injected over the sensor surface at a flow rate of 30 μL/min at 25 °C with an association time of 60 s and a dissociation time of 60 s. Solvent corrections were performed using running buffers containing 1.6%–2.4% DMSO. Data were fitted with steady-state affinity analysis using Biacore 8K evaluation software.

### Crystallization and Fragment Soaking

Crystallization was performed at 4 °C using the hanging-drop vapor diffusion method. The reservoir contained 1.6 M sodium malonate (pH 6.0). Fragment soaking was conducted by mixing 0.5 µL of 100 mM fragment solution (in ddH₂O) with 0.5 µL reservoir solution and adding the mixture to the crystallization drop. The final fragment concentration was approximately 20 mM. Crystals were soaked for 24 hours, cryoprotected in reservoir buffer containing 25% glycerol, and flash-cooled in liquid nitrogen.

### Data Collection and Structure Determination

Diffraction data were collected at beamlines BL19U1^77^. Data were processed using HKL3000^78^ followed by Aimless from the CCP4 suite^79^. Initial phases were determined using Phenix.Phaser^80^, with PDB entry 7OFU as the search model. Model building and refinement were performed iteratively using Coot^81^ and Phenix.refine^80^. Data collection and refinement statistics are summarized in Table S1.

## Data Availability

The benchmarking datasets, peptide dataset for screening, and fragment database along with for similarity search, along with their results, could be found in Supplementary Information and Supplementary Data. Protein structures in the datasets were downloaded from Protein Data Bank (https://www.rcsb.org/). Information of allosteric sites were fetched from the Allosteric Database (https://mdl.shsmu.edu.cn/ASD/module/mainpage/mainpage.jsp). DUD dataset was downloaded from https://dud.docking.org/jahn/.

## Code Availability

Codes for generating matrixes in CIM format (Ising matrix), CPU-based methods along with test cases are currently available at https://drive.google.com/file/d/1MTKUXVQqYrEBDwo6zNWuh_-M1cm8bLPq/view?usp=sharing.

Construction of Ising matrixes were realized with ProDy 2.4.1 (http://www.bahargroup.org/prody/), NumPy 1.22.2 (https://numpy.org/), sklearn 1.3.2 (https://scikit-learn.org/stable/index.html), RdKit (https://www.rdkit.org/docs/index.html) and kaiwu SDK (in the package). Note that this SDK only work with Python 3.8 (https://www.python.org/). The CPU-based methods were realized with the same packages and FPocket 2 (https://fpocket.sourceforge.net/), and were accelerated by numba (https://github.com/numba/numba). Scoring of peptides were realized in a Python 3.9 environment, with openmm 7.7.0 (https://openmm.org/) and cg2all (https://github.com/huhlim/cg2all).

Data analysis were done with Python 3.8, NumPy 1.22.2, SciPy 1.10.1 (https://scipy.org/), Pandas 2.0.3 (https://pandas.pydata.org/) and Matplotlib 3.7.3 (https://matplotlib.org/).

## Acknowledgements

This study was partly supported by grants from the National Key R&D program of China (2023YFF1205103 to J.Zhang), the National Natural Science Foundation of China (81925034, 82441035, 22237005 to J.Zhang, 32300531 to X.L.), Innovative research team of high-level local universities in Shanghai (SHSMU-ZDCX20212700 to J.Zhang), the Starry Night Science Fund of Zhejiang University Shanghai Institute for Advanced Study (SN-ZJU-SIAS-007 to J.Zhang.), the Key Research and Development Program of Ningxia Hui Autonomous Region (2022BEG01002 to J.Zhang), Shanghai Municipal Health Commission (2025ZHYL038 to J. Zhang), and Lingang Laboratory (LG8888 to J. Zhang)

## Author Contributions

J.Zhang and K.W. conceptualized the study. J.Zha developed the encoding algorithms of allosteric-site detection and protein-peptide docking. S.C., and C.C. developed the algorithm of intermolecular similarity calculation. L.Y., and Q.G. developed the coherent Ising machine and performed tests of the hardware. J.Zha, S.C., M.L. and C.W. performed computational experiments. X.Lan performed experiments of GALR1. Y.L., S.N. and C.W. performed experiments of SIRT2. J.Zhong, H.Q., X.Y., Yao Ma and Z.A. performed experiments of APC-Asef. X.S., M.L., Y.Z, H.Q and H.H. performed experiments of NTSR1. J.Zhong performed experiments of AURKA, CDK2 and PLPRO. Y.Cui and Y.Chen performed experiments of SIRT3. W.Q., Q.X. and T.W. solved the crystal structure of PLPRO-Frag160 complex. J.Zha, S.C. and P.G. did the visualization. J.Zhang, K.W., Yin Ma, H.W., Q.G., C.C., Z.A., X.Liu. and Z.Z. supervised the study. J.Zha, S.C., L.Y., J.Zhong wrote the original draft. All the authors reviewed and edited the article.

## Extended Data

**Extended Data Fig. 1.**
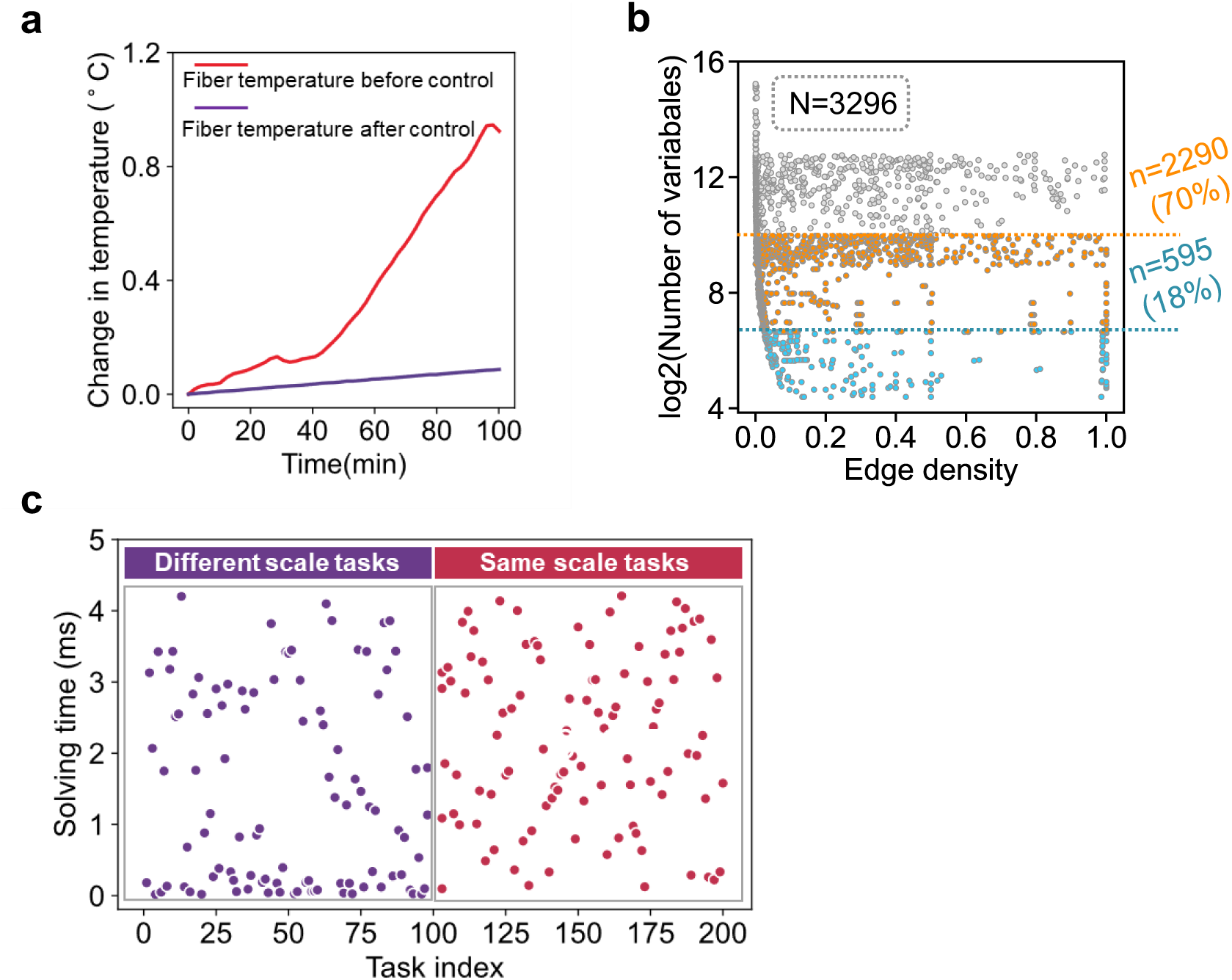
**a.** Temperature change of optical fiber before and after temperature control. **b.** The coverage of COPs that can be solved by different scale hardware, blue dots indicate COPs within 100 bits, representing 18% of the dataset, yellow dots indicate within 2000 bits, representing 70% of the dataset. **c.** Solving time distribution of 100 tasks of different scales (purple dots) and the same scale tasks calculate 100 times (red dots).

**Extended Data Fig. 2.**
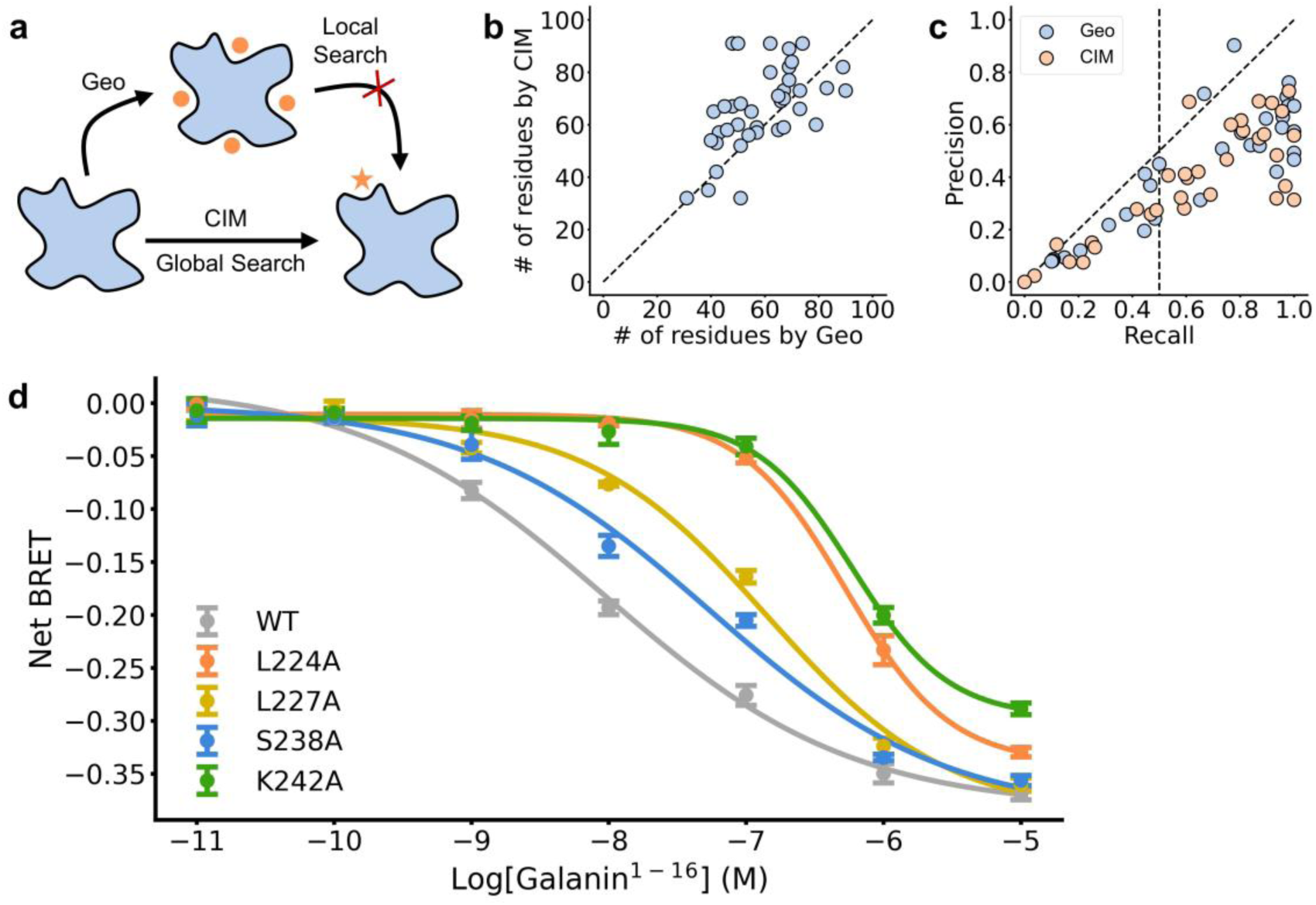
**a.** Temperature change of optical fiber before and after temperature control. **b.** The coverage of COPs that can be solved by different scale hardware, blue dots indicate COPs within 100 bits, representing 18% of the dataset, yellow dots indicate within 2000 bits, representing 70% of the dataset. **c.** Solving time distribution of 100 tasks of different scales (purple dots) and the same scale tasks calculate 100 times (red dots). **d.** Dose-depended response of galanin to different mutant of GALR1.

**Extended Data Fig. 3.**
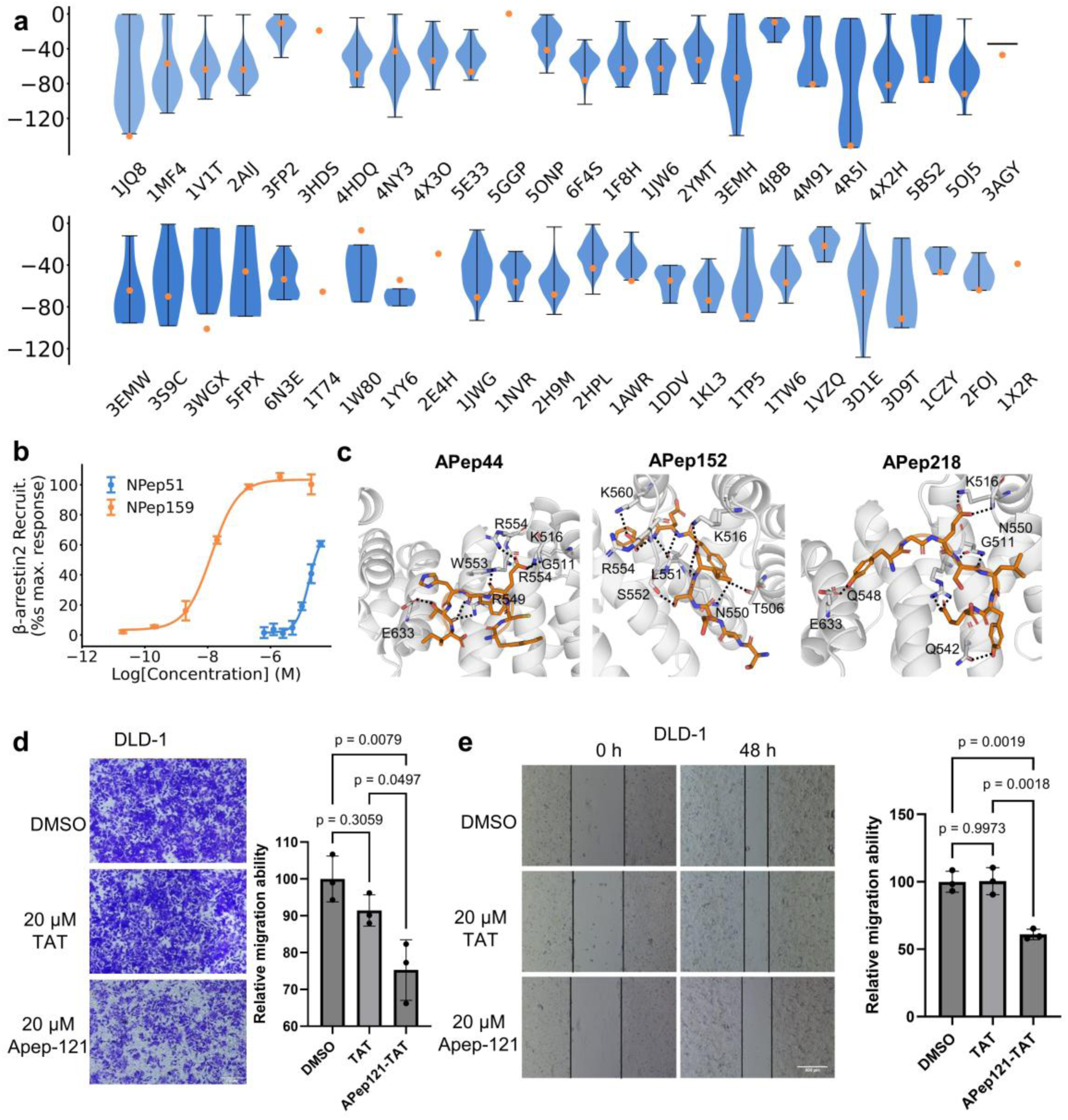
**a.** Distribution of the sampling score by CIM (violin plot) and the sampling score of the crystal structure (orange dot). Dose-depend response curve of the selected peptides**. c.** Docking pose of selected APC-Asef inhibitors. **d.** Transwell migration assays (DLD-1) of APep121-TAT, where TAT is the cell-penetrating peptide. **e.** Wound healing assays (DLD-1) of APep121-TAT. N=3 for all experiments. The error bars show the standard deviation.

**Extended Data Fig. 4.**
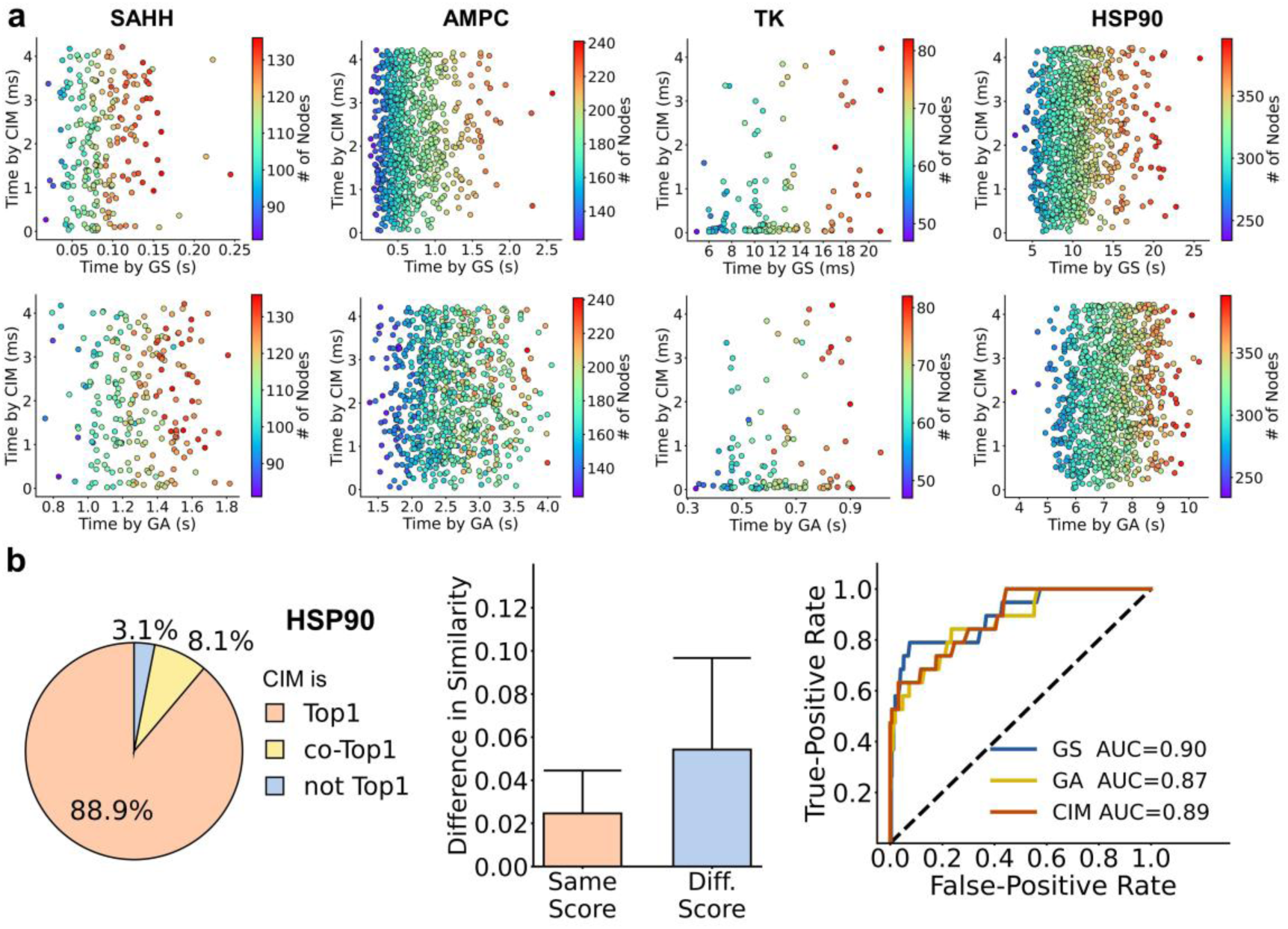
**a**, Comparison of the solving time by coherent Ising machine (CIM), greedy search (GS) and genetic algorithm (GA). **b.** Benchmarking CIM-based similarity calculation on HSP90 and comparison with GS and GA. Left: Quality of solved max-weighted independent set (MWIS). Middle: Influence of calculated similarity by MWIS score. Right: Receiver operating curve (ROC) by the calculated similarity in classifications of molecules.

**Extended Data Fig. 5.**
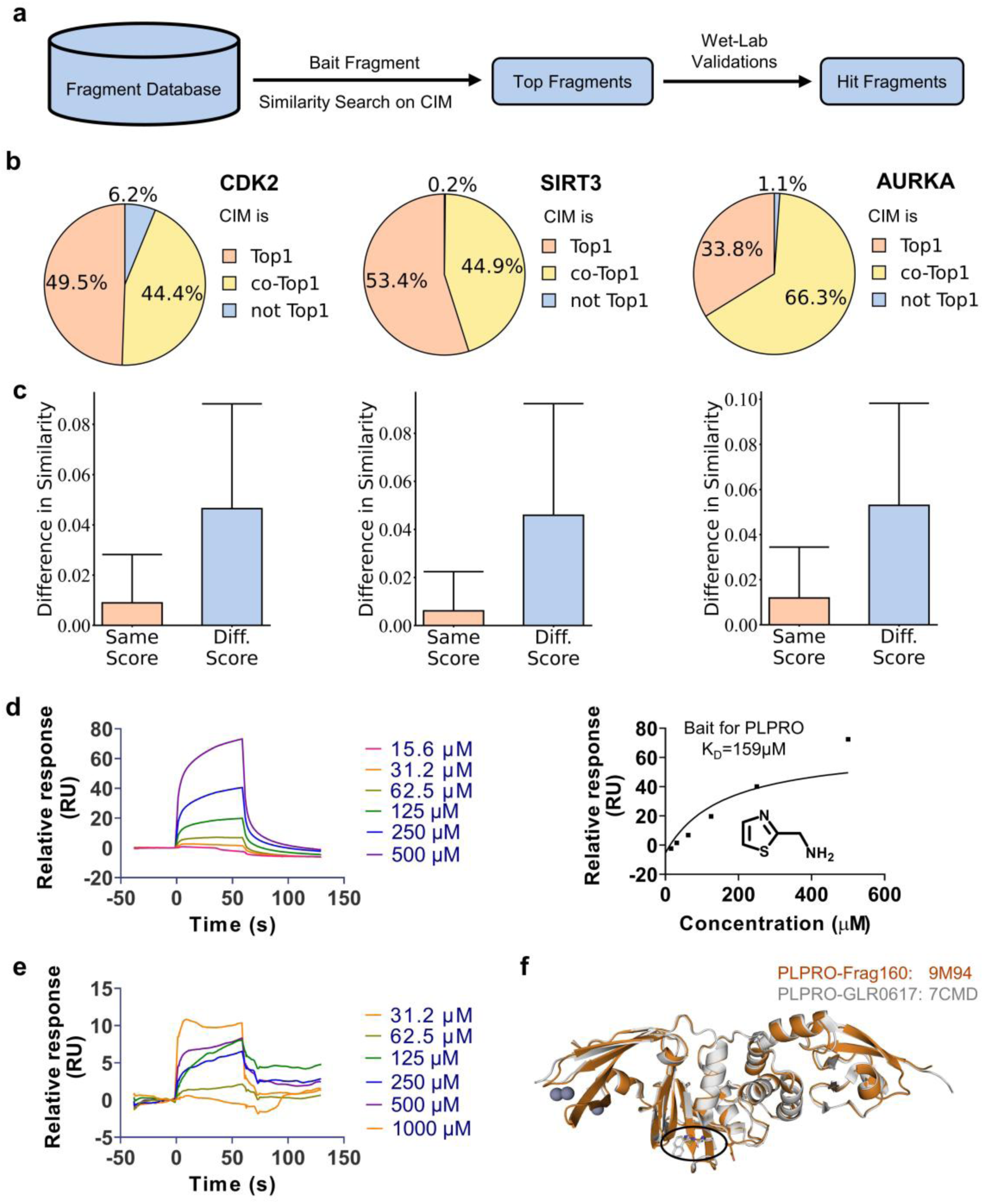
**a.** Workflow of ligand-based drug discovery by CIM. **b-c.** Quality of solved max-weighted independent set (MWIS) (**b**) and the influence of calculated similarity (**c**) by MWIS score in AURKA, SIRT3 and CDK2. **d.** SPR experiment of the bait for PLPRO. **e.** SPR experiment of the Frag160. **f.** Crystal structure of PLPRO-Frag160 complex compared to PLPRO-GLR0617 complex.

